# Immunological female role tested on artificial plugs in three scorpion species

**DOI:** 10.1101/479485

**Authors:** Mariela A. Oviedo-Diego, Camilo I. Mattoni, Alfredo V. Peretti

**Author notes:** Corresponding author E-mail address (MO).

## Abstract

Within arachnids, genital plugs are morphologically diverse, and they can be formed by male, female or be a contribution of both sexes. Although several species of scorpions with genital plugs are known, the physiological effects on the female after being plugged have not been well studied yet. This work compares three scorpion species, two with genital plugs and one without. We first describe the genital plugs morphology of two *Urophonius* species. Second, through the placement of artificial genital plugs in the female genital atrium, we tested 1) whether there are interspecific differences in the immune encapsulation response on the artificial genital plug, 2) if there are an effect in the hemocyte load in the hemolymph, and 3) if individual’s immunological parameters and body weight are correlated. Additionally, we describe and quantify the hemocytes in these species. In both species of *Urophonius*, genital plugs were found covering the female genital aperture and blocking the genital atrium. The plugs consist of three zones that are distinct in morphology and coloration. We found different patterns of encapsulation and melanization on the artificial plug according to the species, with a greater and more specific response in females of plug producing species. Also, these species showed a decrease in the hemocyte load one month after the placement of the artificial plug, possibly due to the recirculation of the hemocytes into the genital area. In addition, correlations were found between the body weight and the immunological parameters, as well as between different immunological parameters. Our results suggest that females contribute to the formation of genital plugs by adding material and generating the darkening of the genital plugs in certain zones. This comparative study can help to provide a wider framework of different physiological consequences related to a particular postcopulatory mechanism such as the genital plugs.

## Introduction

Among the many reproductive strategies that organisms exhibit, there are some that involve males’ adaptations favored by sperm competition to increase their reproductive success [1-3]. Males would compete for the monopolization of females toward preventing, reducing or avoiding sperm competition [2,4]. Genital plugs are structures that block or cover some portion of the female genitalia after mating, and consequently can prevent sperm competition, acting as mechanical or visual impairments [5-6] or by a physiological or behavioral alteration, such as decreased female receptivity [7-10]. The plugging of females is a widespread phenomenon in the animal kingdom, including insects and arachnids [4, 11-14]. Within arachnids, genital plugs are morphologically diverse, varying according to the taxa. In general, genital plugs can be formed by the coagulation of the male’s ejaculate or glandular substances [12, 15-21], portions of the spermatophores [5,9,11] or even parts of the male’s body or genitalia [22-24]. However, because the plugs could represent the result of a sexual conflict in the domain of fertilization [25-26], sexually antagonistic coevolution would favor counter-adaptations of the females. For example, females can prevent the placement of a plug [27-29], controlling the duration of the mating by means of the formation of a plug [30-31] or actively removing it or degrading it [32-37]. Female control of the fate of the genital plug has been proposed as a mechanism of cryptic female choice [38-39]. This would imply their cooperation (or not) in forming the genital plug, depending on such characteristics of males (e.g., copulatory courtship) [40-41] or characteristics of the genital plug (e.g., the quality of nutritive substances) [42-45] or as a mechanical obstruction to potentially harmful or unnecessary new copulas [46].

The post-mating physiological consequences that are triggered in the female after the deposition of the male’s genital plug have not been well studied. Some studies have described secretions of the epithelium of female genitalia that adhere material to the plug and may help to anchor or consolidate the genital plug, or conversely, this material may degrade the male’s plug [15, 47-48]. In *Drosophila nasuta*, the male transfers a substance within the ejaculate that would activate the phenoloxidase pathway, a humoral component of the immune system on arthropods, and lead to the formation of a large, opaque mass in the female’s uterus [49-50]. In other cases, females produce proteases that degrade the genital plug and, in turn, the male’s plug has ejaculatory proteins with specific inhibitors for these proteases, evidencing antagonistic coevolution between males and females on the effectiveness of the genital plug [48]. Several studies have found changes in the immune system after mating [51]. For example, it has been found that, after mating, several immunological parameters may be improved [52] or weakened [53-55]. Also, mating may cause the activation of immune system molecules in reproductive tissues [56-57] or changes in the expression of immunity genes [58].

Arthropods have a relatively simpler immune system than vertebrates, since they lack acquired immunity [59, but see 60], although this does not mean that the immune system is less specific [61-63]. Immune responses comprise cellular-like responses mediated by the hemocytes-granulocytes (GRs) and plasmatocytes (PLs)-(e.g., coagulation, phagocytosis, nodule formation, encapsulation), and humoral-mediated responses (e.g., complement-like proteins, antimicrobial peptides, products generated by the phenoloxidase pathway) [64-66]. In particular, the encapsulation response (i.e., hemocytes’ adhesion in tight layers around an extrinsic factor) involves the action of GRs that recognize the extrinsic factor and release granules (with chemical signals of recruitment of PLs, enzymes and precursors for melanin synthesis and ‘encapsulation-promoting factors’) [64,67]. Therefore, the capsule formation is associated with melanization produced by the prophenoloxidase (proPO) cascade (activation of the phenoloxidase enzyme) with reactive oxygen and nitrogen species emitted and targeted against the extrinsic factor [68]. In chelicerates, some studies investigate hemocyte ultrastructure [69-70] and the presence of antimicrobial molecules in the hemolymph [71-73]. But we still need studies that evaluate the relationship between postcopulatory mechanisms and immune response parameters. In the framework of the theory of immunocompetence, higher quality individuals are better able to meet the costs of maintaining good sexual characters and good immunological defense, and will therefore be preferred as couples [74-75].

The reproductive biology of scorpions has certain characteristics that make it a potentially useful model for the study of these topics. Their courtship is complex and ritualized, after which males adhere a sclerotized spermatophore to the soil, from which the female receives the sperm [76-78]. The sperm penetrates the genital aperture of female and advances through the genital atrium towards the seminal receptacles [79]. After fertilization, the viviparous embryos develop within the ovariuterus until the time of parturition [80-81]. In many species of scorpions, females present a genital plug after the sperm transfer [12], although their morphology is very diverse and the function is discussed [11]. The efficacy in preventing sperm competition is in many cases linked to the morphology of the genital plug. It has been observed that in *Mesomexovis punctatus* (Karsch, 1879) inseminated females experience a decrease in sexual receptivity, that coincides with the presence of plugs with strong anchoring mechanisms to the genital atrium and obstruction of the genital aperture [9,19]. In contrast, in *Bothriurus bonariensis* (C.L Koch, 1842) this fall in receptivity after mating is not observed, coinciding with a small and membranous plug that does not block the female genital aperture [82]. However, a strict relationship between plug size and efficiency in the sperm competition avoidance should be reviewed in depth in scorpions, since other factors might affect the effectiveness of the plug (e.g., accompanying chemical substances, anchoring and blocking mechanisms, hormonal activator-deactivators, triggering factors of the immune response). In some cases, the formation of the genital plug is almost completely attributed to the male [9,12,19], although the participation of the female has been suggested [11-12,19,83-84]. In ultrastructural studies of the atrial epithelium of the female, pores and glandular cells and high secretory activity have been described, so the possibility of female participation is strongly expected [18,84-85].

In this work, females of three scorpion species are compared. *Urophonius brachycentrus* (Thorell, 1877) and *U. achalensis* (Ábalos and Hominal, 1974) [11,86] two species of the family Bothriuridae that have genital plugs were compred with *Zabius fuscus* (Thorell 1877), a buthid species with no genital plug [12]. A wide variety of genital plugs are known in Bothriuridae, which may be ‘gel-like’ (formed by sperm or accessory gland substances) [11,18,82] or ‘sclerotized’ plugs (derived from cuticular portions that detach from the spermatophore) [11,18]. Within sclerotized plugs, there are two subtypes: simple (filamental, membranous) and complex plugs (cone-shaped, mixed) [12]. It has been proposed that *Urophonius* has a ‘mixed’ plug since it presents a combination of detachable portions of the spermatophore and glandular substances [11]. It is known that males of *Urophonius brachycentrus* and *U. achalensis* transfer an ‘initial plug’ formed by two hemi-mating plugs (one per hemispermatophore) that join in the formation of the spermatophore during sperm transfer. This ‘initial plug’ presents a translucent coloration when it has just been transferred to the female and shows a progressive darkening during the reproductive season (see below the results of this study). Certain changes in the size and coloration of the plugs of *Urophonius*, and other species [9,11,19], might be linked to immunological responses such as encapsulation and melanization. These reactions strongly resemble the immune response that is activated on artificial implants (e.g., nylon filaments) that are inserted into the hemocoel of individuals and that, after a time, present areas with dark encapsulations [87-90].

To elucidate these questions about the female role in the formation of genital plugs, we first described the genital plugs and their positioning within the female genital atrium in both *Urophonius* species. In second place we evaluated the encapsulation immune response to a mechanical stimulus, similar to that of the genital plug, in the female genital atrium. We expected to find larger, dark-colored (melanotic) encapsulations in the artificial genital plugs of females of species that have a genital plug, resembling those observed in the true genital plugs of these species. We used this experimental approach due to the impossibility of replacing extracted plugs in the females since these are extremely fragile and strongly anchored within the female genital atrium (Oviedo-Diego, Mattoni and Peretti, unpublished data). Thirdly, we surveyed the types of hemocytes present in the hemolymph and total hemocyte load (THL) of the females of each species. It was observed if there were changes in the THL before and after the artificial plug placement. We expected recirculation of the hemocytes toward the genital area if these are involved in the changes observed in the artificial plugs. Finally, we looked for relations between the immunological parameters (encapsulation areas/coloration as a proxy of melanization and THL) and between the parameters and the body weight of individuals, since heavier females could have greater THL and be more competent to face an immune challenge such as the artificial genital plug.

## Materials and methods

### Studied species, collection and rearing

Two sister species of the family Bothriuridae were studied. *Urophonius brachycentrus* and *U. achalensis* present winter surface activity [91] and were collected at the beginning of the season (from May to June). The time of collection was determined to ensure that the females were virgins and did not have a genital plug, as when the females are inseminated they always present a genital plug, and their distal portion is visible below the genital operculum (Fig 1A). In contrast, *Zabius fuscus* individuals are active in summer (November to March) [92] and inseminated females do not present genital plug [12,18]. Individuals were collected during the day by turning rocks over in the Sierras Grandes at altitudes from 800 to 1900 MASL. Although *Zabius fuscus* (Fam. Buthidae) is phylogenetically distant from the two species of Bothriuridae, it was chosen for this study because as far as we know there are no bothriurid species that do not have a genital plug [11-12,18,82,93-94]. In the laboratory, each specimen was weighed with a digital balance (Ohaus Pioneer PA114). The scorpions were conditioned in individual plastic containers (9 cm x 6 cm) and were kept with moistened cotton as a water supply, and fed once a week with larvae of *Tenebrio molitor* (Coleoptera, Tenebrionidae) or adults of *Shelfordella tartara* (Blattodea, Blattidae). The specimens were maintained at constant temperatures (10°C in winter, 25°C in summer). Voucher specimens were deposited in the collection of Laboratorio de Biología Reproductiva y Evolución, Universidad Nacional de Córdoba, Argentina.

**Fig 1.**
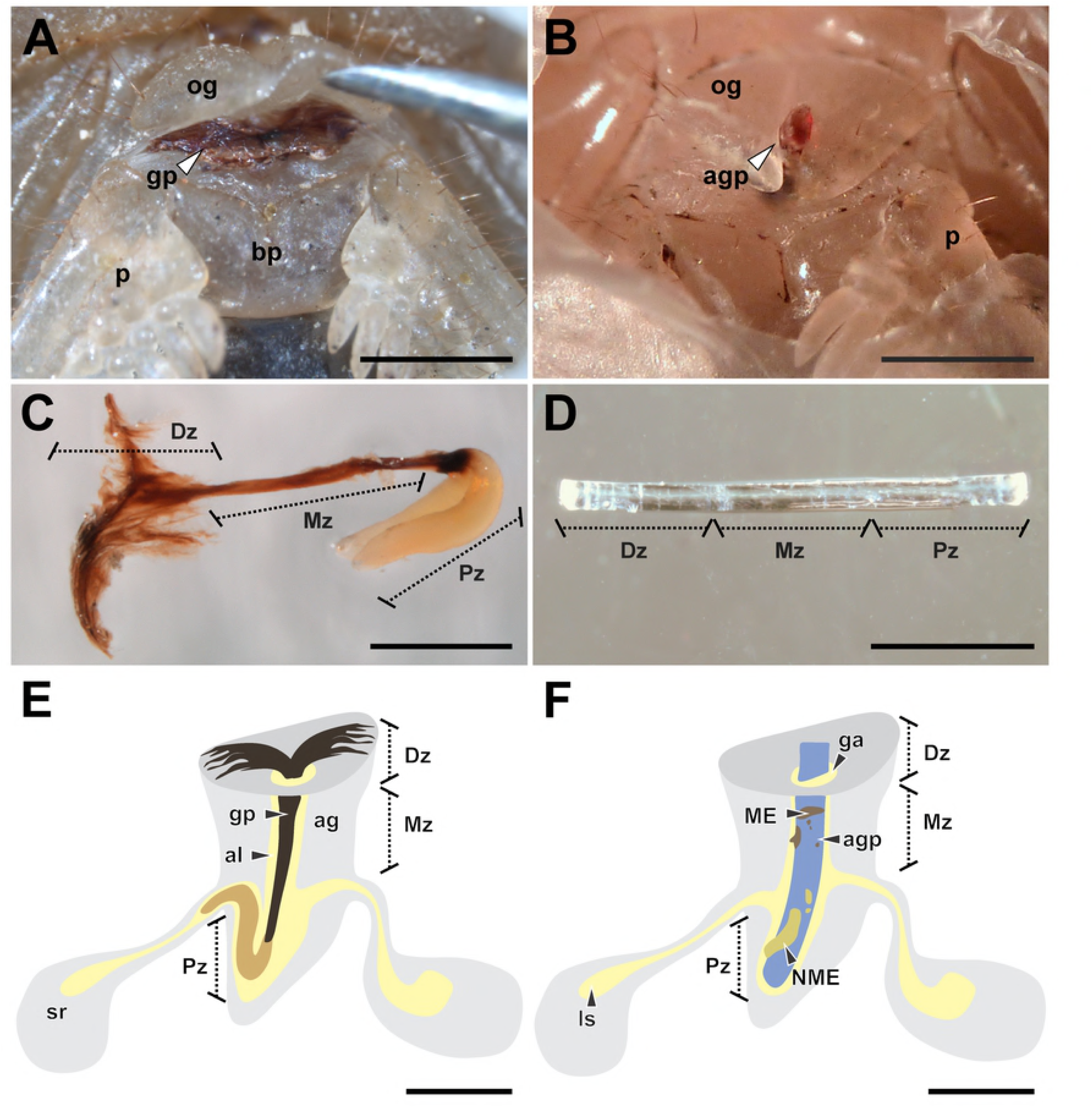
Genital plugs and artificial genital plugs of the study species. (A) ‘Distal’ zone of genital plug below the female genital operculum of *Urophonius achalensis*. (B) Protruding distal portion of the artificial genital plug positioned within the female genital atrium of *Urophonius achalensis*. (C) Genital plug extracted of an inseminated female of *Urophonius brachycentrus*.(D) Artificial genital plug before being placed on a female. (E) Scheme of a genital plug (*Urophonius*) and its positioning within the female genital atrium. (F) Scheme of an artificial genital plug and its positioning within the female genital atrium. Abbreviations: ag, genital atrium; agp, artificial genital plug; al, lumen of the genital atrium; bp, basal piece; Dz, distal zone; ga, genital aperture; gp, genital plug; ls, lumen of the seminal receptacle; ME, melanotic encapsulation; Mz, middle zone; NME, non-melanotic encapsulation; og, genital operculum; p, pectine; Pz, proximal zone; sr, seminal receptacle. Scale bars: 1 mm.

### Morphology of genital plugs and positioning within female

To observe the positioning of the plug within inseminated females (*U. brachycentrus* N=20; *U. achalensis* N=20) dissections were performed. The specimens were sacrificed in a freezer at −20 ° C for fifteen minutes and then dissected under a stereoscopic microscope (Nikon SMZ 1500). After the dissection, the genital plug was removed from the female’s atrium with straight tweezers. The dissected specimens and genital plugs were photographed with a digital camera (Nikon Digital Sight DS-FI1-U2) coupled to the stereoscopic microscope. Some genital plugs (N=10 per species) were kept in a 1 mL microcentrifuge tube exposed to the air and were photographed every week for a month. In this way, we observed if there were changes in the coloration or morphology of the genital plugs outside the female (e.g., by oxidation of the plug material). We also evaluated changes in the coloration and morphology of the plugs inside the females throughout the reproductive season (N=10 per species) by examining the external portion of the plugs below the genital operculum.

### Encapsulation response on the artificial genital plug

#### Placement and removal of artificial plug

A piece of sterile nylon monofilament (3 mm x 0.1 mm) was placed in the genital atrium of each female, resembling in size and positioning to the genital plug present in females of the species of *Urophonius* (Fig 1). The specimens were immobilized on a microscope slide with Parafilm®. A small hole was made in the parafilm to access the genital operculum of the individual, and the artificial plug was gently inserted up to the end of the atrium by lifting the operculum with straight tweezers (Fig 1B, D, F). The surface of the artificial plug was slightly roughened with sandpaper to reach a rough surface and enhance the adhesion of hemocytes to the artificial genital plugs [95-96]. This procedure does not cause any damage to the female genitalia. The artificial plug was left in the genital atrium of the female for a month, since it has been observed that the genital plug of *Urophonius* takes approximately this time to present some darkening (Oviedo-Diego, Mattoni, Peretti personal observations). After this period, the artificial plugs were carefully removed, and the tissue remnants were cleaned. Each artificial plug was photographed from two perspectives (front and back), rotating 180º [97] with a digital camera (Nikon Digital Sight DS-FI1-U2) coupled to a stereomicroscope (Nikon SMZ1500). A photographic protocol was used that kept light exposure and magnification constant. Then the artificial genital plugs were preserved in ethanol 80%.

#### Area and coloration of the encapsulations on the artificial plug

The encapsulations on each artificial plug were measured by processing the images with ImageJ 1.45 software [98]. For statistical analysis, the artificial plugs were divided into three zones, since it is known that the genital plugs of the *Urophonius* species studied also have three zones (‘distal’, ‘middle’ and ‘proximal’ to the body of the individual) (See Results). The zones of the artificial plugs were defined by dividing the total length of the filament (3 mm) into three parts of equal length so that each zone was 1 mm long. The area that remained within the body of the female contacting the end of the genital atrium was the ‘proximal’ zone of the artificial plug, while the zone more distal to the body of female was the ‘distal’ zone of the artificial plug. The areas of the encapsulations were compared between the zones of the artificial plugs and between species. The encapsulations were classified as melanotic (ME) or non-melanotic (NME) according to their coloration (Fig 1F). The coloration was calculated with the average grayscale value from the pixels of the different areas of the artificial plug encapsulations. The 0 value represents black and 255, white. The classification of encapsulations coloration was carried out using a threshold value of 50 in the average grayscale, being ME if the color was lower than the threshold value and NME if it was higher than this value.

### Extraction, characterization and quantification of hemocytes

The second left leg of individuals was completely excised (between tarsus and tibia) to allow a considerable drop of hemolymph to flow from the wound. A sample of 0.75 μl of hemolymph was taken with a glass capillary from the wound [99]. This sample was mixed with 9.25 μl of Spider Saline Solution [100] in a microcentrifuge tube. A five-second pulse of vortex was carried out three times to the solution to homogenize the sample. Immediately after, the sample was placed in a Neubauer chamber for counting under a light microscope with a phase contrast objective 100x (Nikon Eclipse 50i) [101] at 400X. All hemocytes from virgin females of the different species were identified and counted [102]. The characterization of hemocytes was performed by observing and photographing their characteristics with a digital camera coupled to the microscope (Nikon Digital Sight DS-FI1-U2). The total hemocyte load (THL) (number of hemocytes per milliliter of hemolymph) was compared in two stages: before the placement, and after extraction of the artificial genital plug (through a second cut of the same leg).

### Statistical analyses

We analyze the data with generalized linear mixed models (GLMM). In the analysis of the artificial plugs encapsulations the variables response were the ME and NME areas (mm^2^) and the average grayscale value of each type of encapsulation. The zone of the artificial plug (‘distal’, ‘middle’ and ‘proximal’), the species and the body weight of the individuals were the fixed effects. The body weight of the individuals was measured before and after the placement of the artificial plugs and since there were no significant differences between both instances (Mann– Whitney U test; Z = 0.409, p = 0.683) the average weight value for statistical analyzes was considered. In the quantification of hemocytes the variable response was the THL, and the fixed factors were the species, the stage of quantification (before and after the placement of the artificial plug) and the body weight of the individuals. We also evaluated the possible interactions between the fixed factors analyzed. The individuals’ identity was included in all the models as a random effect. If the random effect variance was small, the effect of the random variable was discarded. Normality and homogeneity of variances of the variables were assessed graphically and analytically. If the assumptions were not met, the variable according to the best distribution was modeled. The coloration of ME and the THL presented a normal distribution. The areas of ME and NME and the coloration of NME presented a gamma distribution, so they were modeled using the glmmadmb function [103]. We used the package lme4 [104] and lsmeans [105] for a posteriori tests in R v. 3.3.3 64 bit [106]. Also, multiple correlations with the Spearman’s method were performed between the different immunological parameters including the three species of scorpions together, and between these parameters and the individuals’ body weight. A significance level α of 0.05 was considered.

## Results

### Morphology of genital plugs and positioning within female

In both species, the genital plug adjusted exactly to the female’s atrium and blocked the lumen and the genital aperture. The genital plugs presented three double-shaped zones (Fig 1C and 2). The ‘distal zone’ to the individual’s body was visible from the outside, and extended below the genital operculum covering the genital aperture (Fig 1A). It was always sclerotized, brittle and darkly colored. In *U. brachycentrus* this zone resembled two thin ‘wings’. In contrast, in *U. achalensis*, this zone was wider with concave platform shape towards the genital aperture. Next to the ‘distal’ zone was the ‘middle’ zone, also sclerotized and dark, formed by two fused structures running along the lumen of the female atrium. While in *U. brachycentrus* this zone was thin and long, in *U. achalensis* it was shorter and it was sometimes more difficult to distinguish the fused structures. Finally, the ‘proximal’ zone consisted of one or more sacciform globular structures, with a flexible gelatinous consistency and a white-yellowish coloration. Projections ascended from the end of the genital atrium to the duct of one of the spermathecae, sometimes occluding the duct (Fig 1E). In *U. brachycentrus* two projections were always found in the ‘proximal’ zone, while in *U. achalensis* the number was variable from one to four proximal projections (Fig 2). We found that the plug undergoes changes in coloration and morphology over time in the genitalia of the female. After mating the plugs presented the ‘distal’ zone (visible below the operculum) with translucent coloration and a thin, fragile consistency. As the reproductive season progressed, the plug darkened and acquired a sclerotized consistency (June to August). Towards the end of the season before parturition (November to December), a decrease in the size of this zone of the plug was observed. Conversely, no changes were observed in the coloration or morphology of the plugs extracted from the females and exposed to the air.

**Fig 2.**
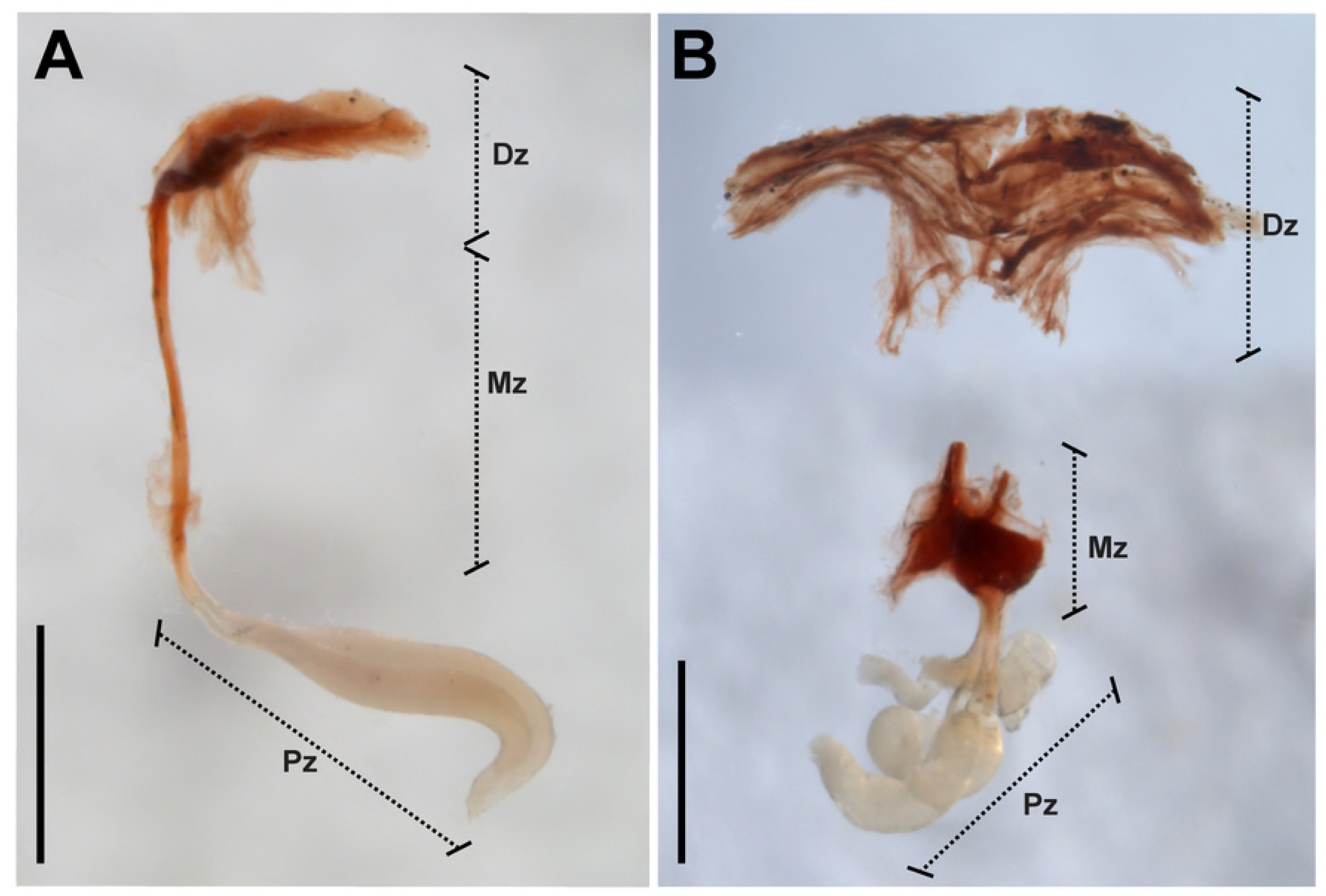
Genital plugs of *Urophonius* species. (A) Genital plug of *Urophonius brachycentrus.* (B) Genital plug of *Urophonius achalensis*, note that the plug has been excised below the distal zone by handling during removal. Abbreviations: Dz, distal zone; Mz, middle zone; Pz, proximal zone. Scale bars: 1 mm.

### Encapsulation on artificial genital plugs

Regarding the encapsulation response of the females, different characteristics and magnitudes of this type of immune response were observed, depending on the species and the zone of the artificial genital plug (Fig 3A-F). Occasionally, a non-melanotic encapsulation response was observed, generally with excrescences of larger surfaces and almost continuously surrounding the genital plug. This type of encapsulation was white-yellowish, translucent or opaque. Melanotic encapsulation presented more specific arrangements, generally in the form of isolated granules in different zones. It was classified according to its color tones from brown to reddish. In all the artificial plugs some type of encapsulation response was found, although in some cases the encapsulations were not present in all the zones of the artificial plug (Fig 3A-F). Hemocytes could be observed on the artificial plugs and in their surroundings under an optical microscope (Fig 4A-B). Sometimes it was possible to see the deposition of a substance around the entrance of the artificial plug by the genital aperture, and even the formation of projections (Fig 4C-D).

**Fig 3.**
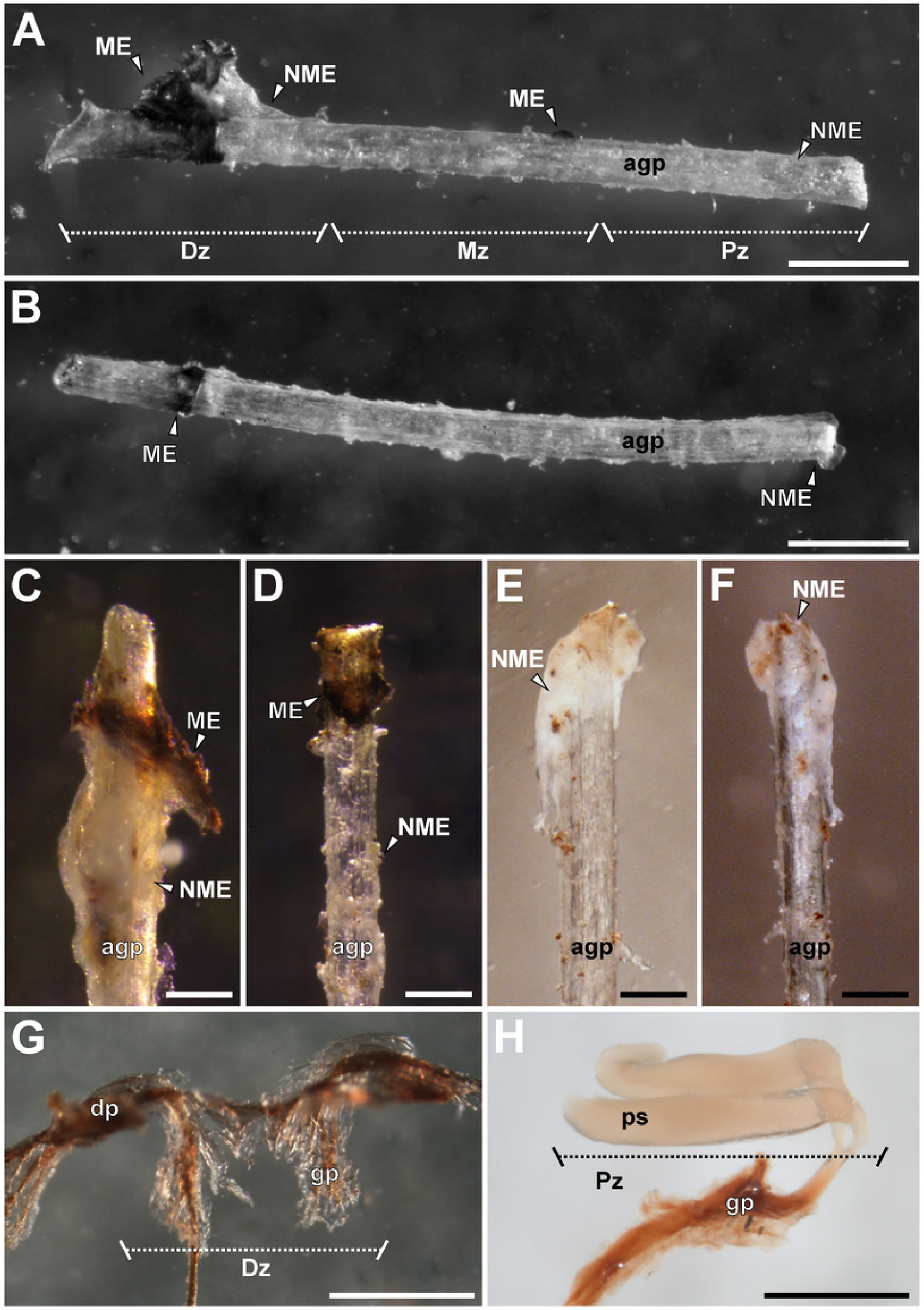
Different encapsulation responses on artificial genital plugs and comparison with *Urophonius* genital plugs. (A) Artificial genital plug with encapsulation response of *Urophonius achalensis.* (B) Artificial genital plug with encapsulation response of *Zabius fuscus.* (C) ‘Distal’ zone of artificial genital plug of *U. achalensis.* (D) ‘Distal’ zone of artificial genital plug of *U. brachycentrus.* (E) ‘Proximal’ zone of artificial genital plug of *U. achalensis.* (F) ‘Proximal’ zone of artificial genital plug of *U. brachycentrus.* (G) ‘Distal’ zone of the genital plug of *U. achalensis.* (H) ‘Proximal’ zone of the genital plug of *U. brachycentrus.* Abbreviations: agp, artificial genital plug; dp, distal projections; Dz, distal zone; gp, genital plug; ME, melanotic encapsulation; Mz, middle zone; NME, non-melanotic encapsulation; ps, proximal sacciform projections; Pz, proximal zone. Scale bars: A, B, G, G= 0.5 mm; C, D, E, F= 0.2 mm.

**Fig 4.**
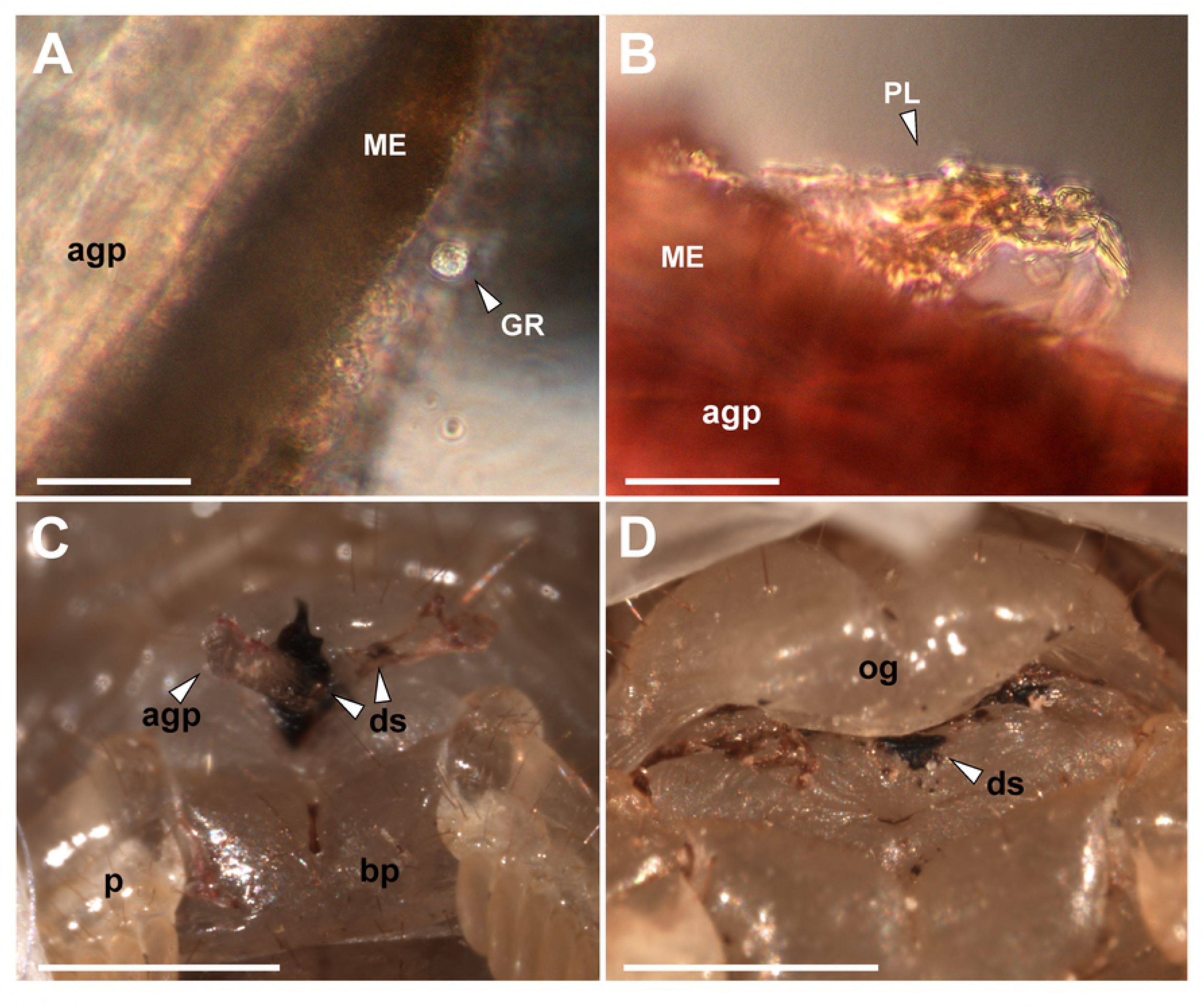
Hemocytes cells and distal substances found on the artificial genital plugs of females of *Urophonius*. (A)(B) Surface of the artificial genital plug of *Urophonius brachycentrus* under an optical microscope, note hemocyte cells. (C)(D) External female genitalia of *Urophonius achalensis*, note artificial genital plug and surrounding substance in the area of the genital aperture. Abbreviations: agp, artificial genital plug; bp, basal piece; ds, distal substance produced by the female; GR, granulocyte; ME, melanotic encapsulation; og, genital operculum; p, pectine; PL, plasmatocyte. Scale bars: A= 0.2 mm; B= 40 μm; C, D=1 mm.

### Areas of encapsulation

We found an effect of the interaction between the species and the encapsulated zone of the artificial plug (ME: Df=2, χ^2^=14.827, p=0.005; NME: Df=2, χ^2^=20.411, p=4.144E-04) (Fig 5A-B and S1 Table). ME values for *U. achalensis* were higher, although they were not significantly different from those of *U. brachycentrus*. In contrast, the two species of *Urophonius* did show differences with the ME values of *Z. fuscus* that were the lowest (Fig 5A). Differences were also found in the areas covered by encapsulations in the different zones of the artificial plug. All species had higher ME in the ‘distal’ zone, although *U. brachycentrus* had fewer differences between zones. As for NME, all species presented similar areas of encapsulations. *Z. fuscus* showed a homogenous, low response in all zones, and *Urophonius* species showed a greater response in the ‘distal’ and ‘proximal’ zone (Fig 5B). The interaction of body weight with the artificial plug area for NME was significant (Df=2, χ^2^=18.454, p=9.835E-05).

**Fig 5.**
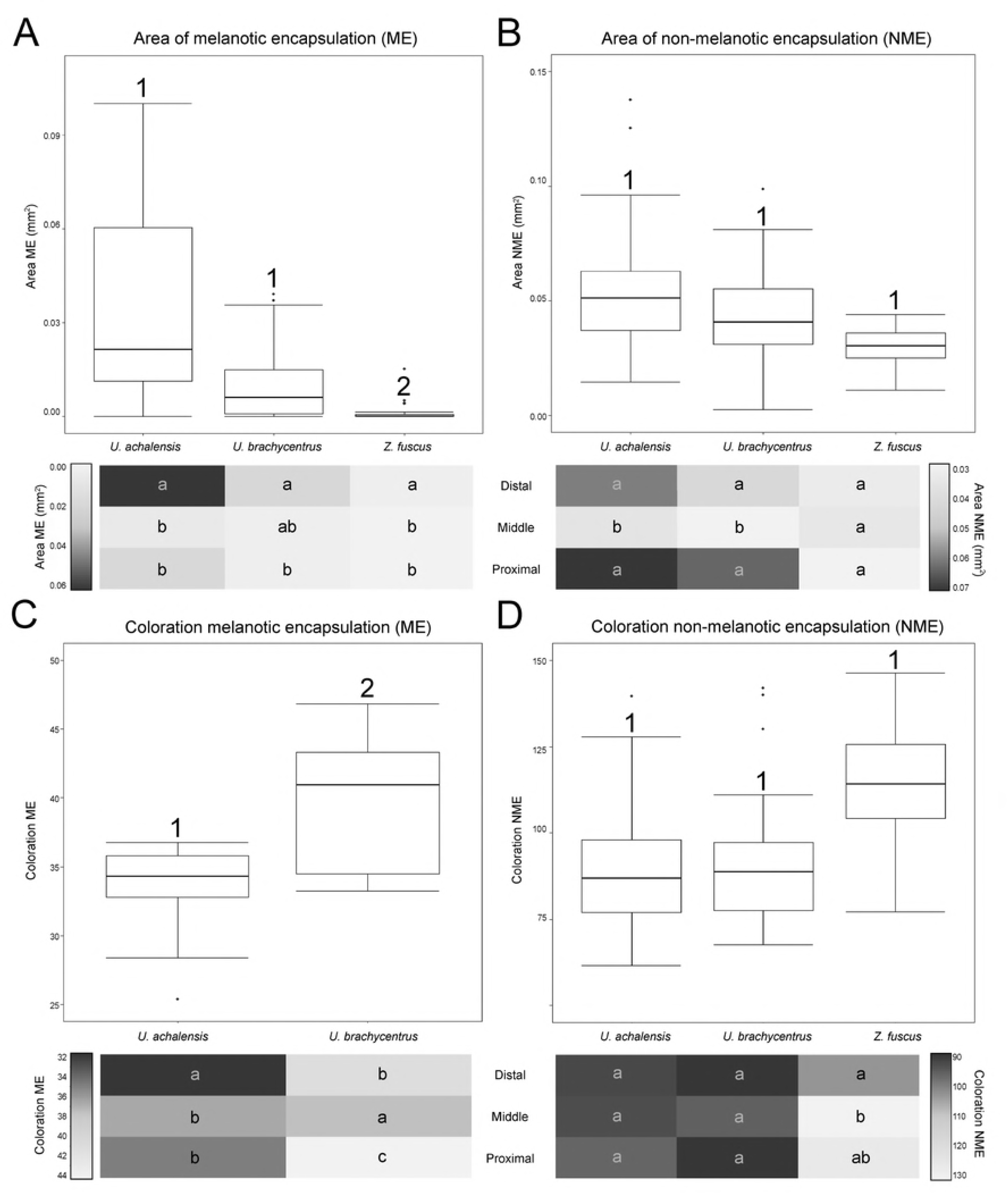
Graphs of immunological parameters of the encapsulation response in females of three scorpion species. Top boxplots showing distribution of data set and differences between species, below heat maps charts in which average values are represented by colors (scale of reference of each variable to the side of the graph) according to the zones of the artificial genital plug. (A) Area of melanotic encapsulation (ME) (mm^2^). (B) Area of non-melanotic encapsulation (NME) (mm^2^). (C) Color of NME encapsulation (average grayscale value). (D) Coloration of ME encapsulation (average grayscale value). Numbers indicate significant differences (p <0.05) between species above the boxplots. Letters on heat maps charts indicate significant differences (p <0.05) between zones of the artificial genital plug within each species (i.e. intraspecific comparison: reading vertically, not horizontally).

### Coloration of the encapsulated zones

Since *Z. fuscus* females rarely presented ME in the ‘middle’ and ‘proximal’ zone of the artificial plugs, only the coloration of these encapsulations was analysed for the females of two *Urophonius* species (Fig 5C and S1 Table). We found a significant interaction between the species and the encapsulated zone (Df=2, χ^2^=29.698, p=3.558E-07). *Urophonius achalensis* females showed lower values on the mean grayscale than *U. brachycentrus*, i.e. darker coloration of ME and there was also variation in the artificial plug zone. *Urophonius achalensis* presented ‘distal’ zones significantly darker than the rest of the zones. In contrast, *U. brachycentrus* presented darker ‘middle’ zones, followed by the ‘distal’ zones and significantly clearer ‘proximal’ zones. For the coloration of NME, we found an effect of the interaction between the species factor and the artificial plug zones (Df=4, χ^2^= 23.463, p=1.023E-04). *Z. fuscus* presented in general clearer encapsulations although they were not significantly different from those of *Urophonius* spp. (Fig 5D and S1 Table). In this species, the ‘distal’ zone was darker than the others. For the *Urophonius* species, all the zones presented NME encapsulations of equal coloration. There was an interaction between body weight and the artificial plug zone for NME (Df=2, χ^2^=10.976, p=0.004).

### Characterization of hemocytes of females of the studied species

We identified different types of hemocyte cells in the hemolymph of the females in the species studied. Hemocytes presented different morphology and size (Fig 6). Granulocytes (GRs) were cells with spherical and isodiametric shapes and regular contours. GRs had cytoplasmic extensions variables in shape, although the extensions were in general short and acute. The cytoplasm of the GRs was dense and always presented abundant refractive oval shaped granules (Fig 6B-C). Plasmatocytes (PLs) were highly variable in shape, generally spindle-shaped with multiple large, rounded cytoplasmic extensions radiating in ameboid form from the central zone. The cytoplasm was hyaline and homogenous, with vacuoles and small or no inclusions (Fig 6E-F). Sometimes the vacuoles occupied a large portion of the cytoplasm of the cell, pushing the nucleus to an eccentric position, giving it a signet-ring appearance with sharp projections (Fig 6F). Also, other types of hemocytes were observed, although they were not found in all the samples. All these hemocytes had a rounded and rather an isodiametric shape, and did not expand cytoplasmic extensions such as PLs. They presented granules in the cytoplasm of different shapes and nature. Cystocytes generally presented a cytoplasm with small granules, a large vacuole and eccentric nucleus. Spherulocytes possessed large, dark granules or spherules of a homogeneous size, which completely obscured the nucleus of the cell. Adipohemocytes presented an eccentric nucleus with typical fat lipid droplets in their cytoplasm. Free cells were observed in the hemolymph and also grouped in agglomerates and, although in these cases it was difficult to identify the clustered cells, we were able to determine that on occasions the clusters may have cells of a different type (Fig 6A, D).

**Fig 6.**
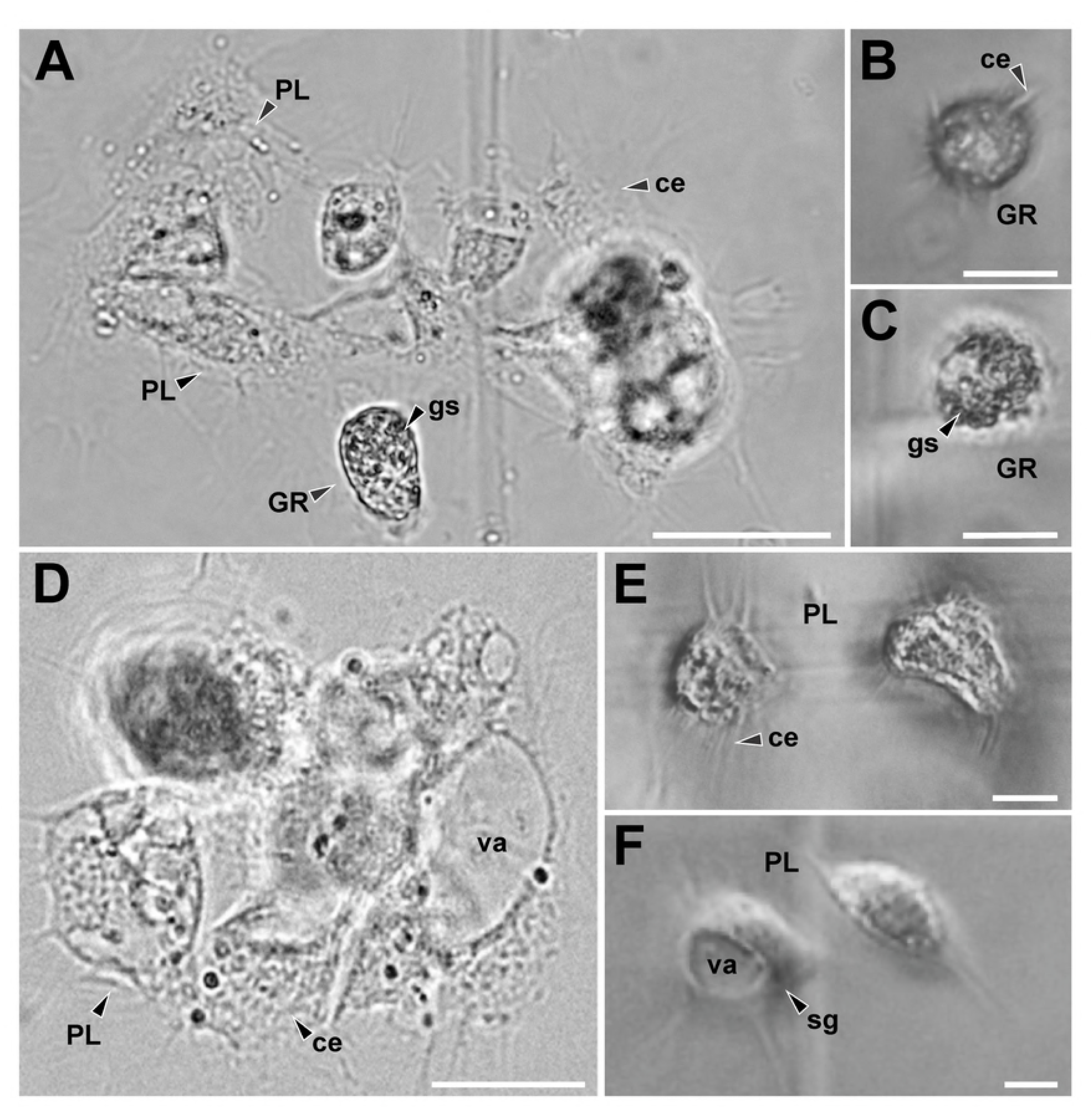
Hemocytes present in the hemolymph of females of three scorpions’ species. (A) Plasmatocytes (PLs) and granulocytes (GRs) cluster of *Urophonius brachycentrus.* (B) GRs of *Urophonius brachycentrus*, note isodiametric morphology, granules in the cytoplasm and short and acute cytoplasmic extensions. (C) GRs of *Zabius fuscus*, note granules in the cytoplasm and short cytoplasmic extensions. (D) PL cluster of *Urophonius achalensis*. (E) PLs of *Zabius fuscus* expanding their cytoplasmic extensions with small inclusions in their cytoplasm. (F) Signet-ring PL of *Urophonius brachycentrus*, note vacuole occupied a large portion of the cytoplasm of the cell. Abbreviations: ce, cytoplasmatic extensions; GR, granulocyte; gs, granules in the cytoplasm; PL, Plasmatocyte; sg, signet-ring plasmatocyte; va, vacuole. Scale bars: A, D= 50 μm; B, C= 20μm; E, F=10 μm.

### Quantification of hemocytes

We found a significant interaction between the fixed factors: ‘species’ and ‘stage of quantification’ (Df=2, χ^2^=35.364, p = 2.093E-08) (Fig 7 and S1 Table). *Zabius fuscus* females showed the highest THL values both before and after the placement of the artificial genital plug with respect to *Urophonius* females. The *Urophonius* spp. showed similar values of THL before the placement of the artificial genital plug. *Zabius fuscus* females showed no difference in THL between the two stages of quantification, while in both species of *Urophonius* females presented a decrease in THL after one month of the placement of the artificial plug, with a greater decrease of *U. brachycentrus* than that of *U. achalensis* (Df=1, F=7.261, p=0.015). Body weight had an effect on THL (Df=1, χ^2^=7.708, p =0.006).

**Fig 7.**
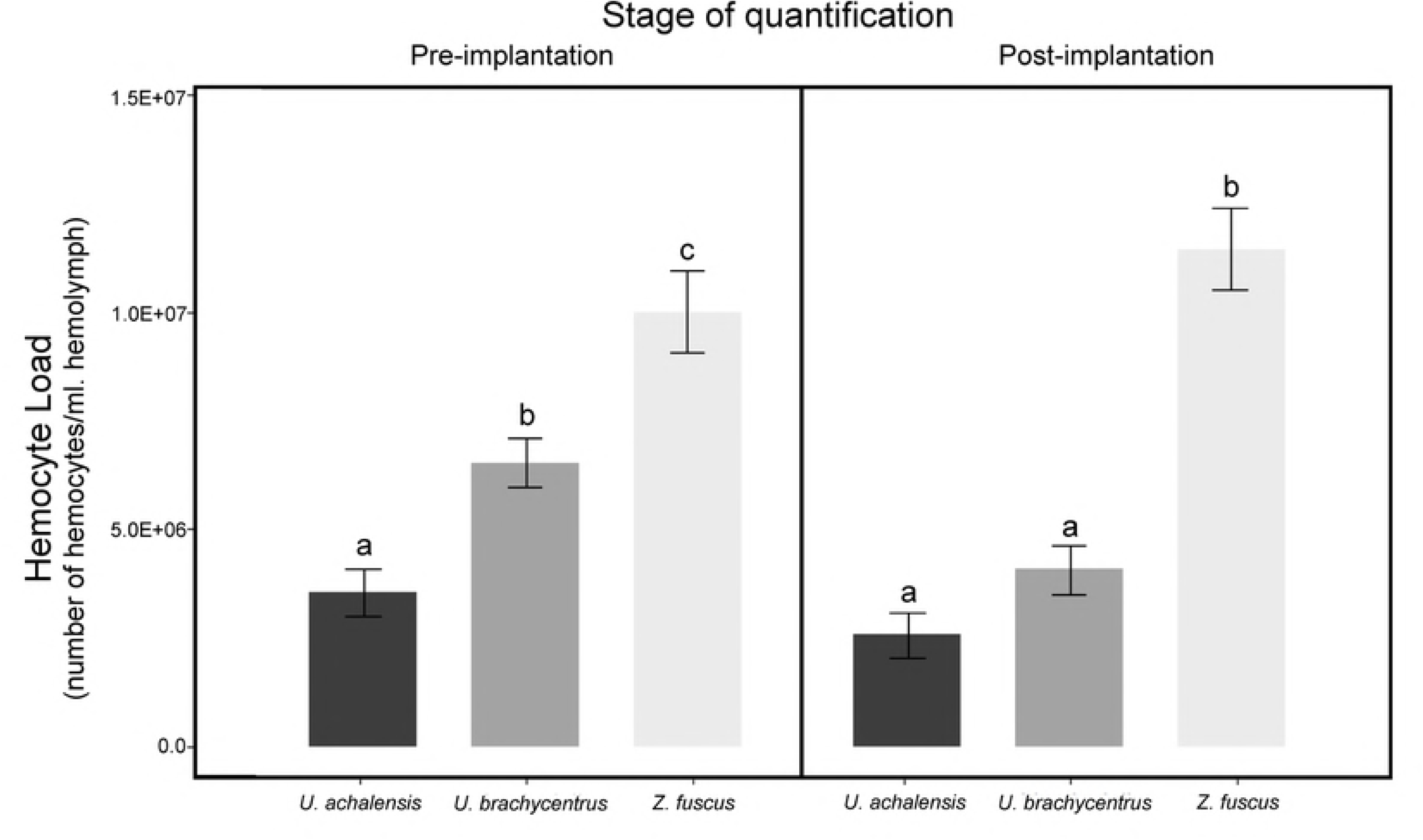
Total Hemocyte Load of females of the three scorpion species. Differences in total hemocyte load (THL) (number of hemocytes per milliliter of hemolymph) according to the stage of quantification (pre-implantation of the artificial genital plug and post-implantation of the artificial genital plug) of three scorpion species. Letters indicate significant differences (p<0.05) between species. Values are shown in scientific E-notation where ‘E’ represents the exponential to 10.

### Correlations between immunological parameters

We noted that the immunological parameters, in general, were highly correlated (Fig 8). A positive correlation was found between the female body weight and the THL but a negative relationship was found between the body weight and the area of both types of encapsulations. Females with a higher THL also presented smaller ME and NME areas on the artificial plugs. However, the THL were positively correlated with the coloration of the encapsulated (i.e. clearer encapsulations). We noted a negative correlation between the area of ME and the coloration of the encapsulated zones, as well as between the NME area and the coloration of the ME encapsulations. A positive correlation was observed between the areas of both types of encapsulation.

**Fig 8.**
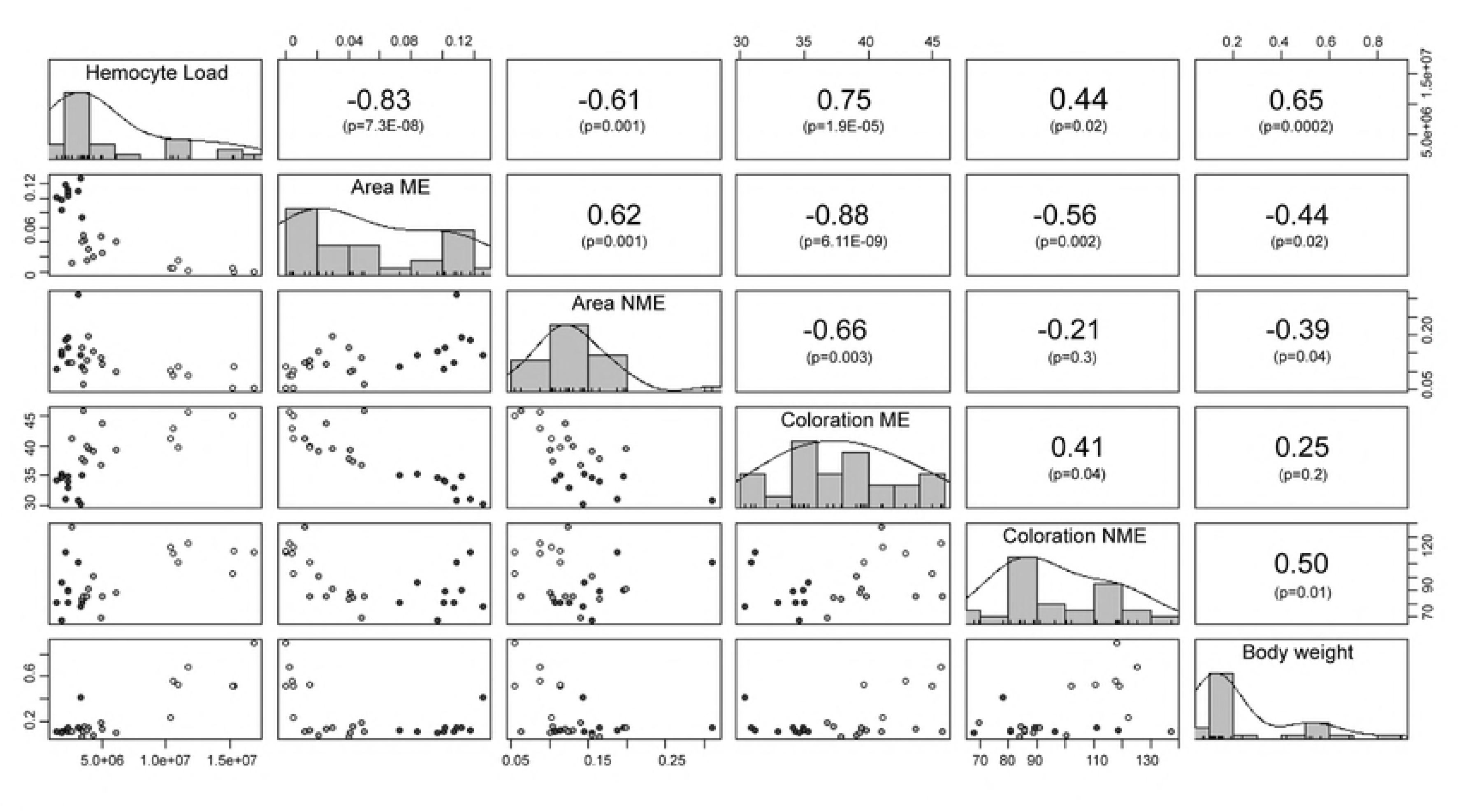
Correlation matrix between the individual’s immunological parameters and body weight. In the diagonal portion of the matrix are found the parameters that were correlated with their corresponding frequency histogram. In the upper portion of the matrix are the Spearman correlation coefficients for each pair of variables and below the correlation p-value. In the lower portion of the matrix are graphs of dispersion of the variables, the colors of points represent the different species of scorpions (Black dots: *Urophonius achalensis*; Grey dots: *Urophonius brachycentrus*; White dots: *Zabius fuscus*). Abbreviations: ME, melanotic encapsulation; NME, non-melanotic encapsulation. Some values are shown in scientific E-notation where ‘E’ represents the exponential to 10.

## Discussion

The description of the morphology of the *U. brachycentrus* and *U. achalensis* genital plugs allowed us to perceive a complex three zoned structure anchored to the female genital atrium. We also observed changes in the plugs’ coloration over time, probably attributable to a female’s role. Given the need for experimental approaches to answer questions about the function or origin of genital plugs [11-13], artificial genital plugs were used in this study to evaluate the female’s contribution to their morphology and coloration. We found that females deposited secretions on the artificial plugs and that some of its zones had darker encapsulations. We confirmed that in plug producing species, the presence of an artificial plug caused a decrease in the hemocyte load. This could mean consequences on the plugged female’s immune system. These results could help to clarify a possible female role in the plugs’ formation and may help to provide a wider framework of different physiological consequences related to this post-copulatory mechanism.

### *Urophonius’* genital plugs and the female’s possible role in its formation

We described inseminated females’ genital plugs of *U. achalensis* and *U. brachycentrus*. In all cases, the plugs blocked completely the lumen of the atrium, the genital aperture, and, in some cases one of the spermathecae’ ducts. Both the female and the male could be involved in the plug’s formation. The double conformation of the plug clearly indicates a male’ s contribution in that, the ‘initial plug’ transferred by the male is formed from the ‘hemi-plugs’ in each hemispermatophore when fused in the spermatophore (involving portions of the ejaculate and glandular products). The changes of the plug over time in morphology, size and coloration could indicate a female’s role. We could discard the hypothesis that these changes of the genital plugs were due to air contact or O_2_ influence. The *Urophonius’* genital plugs present three different zones. Females of *U. achalensis* showed darker melanotic encapsulation in the ‘distal’ zone, and females of *U. brachycentrus* in the ‘middle’ zone. The ‘proximal’ zone of the genital plugs does not present dark coloration, and coincidentally, this was the area in the artificial genital plugs with smaller melanotic encapsulations and clearer coloration. It was also found that the ‘distal’ and ‘proximal’ zones presented the larger non-melanotic encapsulations areas. The projections formed in the ‘distal’ zone of the artificial plugs extending below the genital operculum were similar in shape, size and consistency to those found in the genital plugs. This also adds evidence to the female’s role in the formation of this zone.

In a sexual conflict framework, genital plugs would be strongly associated with the male’s monopolization of females [25-26], and could imply some cost for the females such as a limitation of their possibilities of remating [107]. The plugs that trigger processes, such as those described in this work, could mean a physiological challenge for females, which could in time affect some parameters of fitness or life history. This would be an interesting topic to explore in future experimental studies because the evaluation of these costs could shed light on the discussion of female’s role to the plug formation or degradation.

Another non-exclusive possibility is the occurrence of cryptic female choice [38-39]. This can be manifested as a modulation of the female’s immune response regarding characteristics of the plug (e.g., mechanical effectiveness, chemical composition, size) as well as those of the male (e.g., male’s quality, duration of pre-copulatory and copulatory courtship) [29,40-41,46,108-110]. A comparative and phylogenetic study of the plugs in the Bothriuridae Family could provide information about the evolution of male and female strategies in terms of the plugging phenomenon, and whether this strategy, for example, is related in any way to other traits of the immune system or genital characters.

### An artificial genital plug compromises the immune system of plug producing species

A change in the THL after an immunological challenge has been previously described in species of insects and crabs [111-117]. This change could be explained by the recirculation of free hemocytes in the hemolymph towards affected areas, where the phagocytic or encapsulating action of the hemocytes is necessary. Both the formation of encapsulations on artificial genital plugs and the presence of hemocyte cells in their surroundings evidence a direct action of the hemocytes against this immunological challenge [118]. We found that the encapsulation response varied regarding the analyzed species and the zone of the artificial genital plug. The females of *Z. fuscus* did not show a depletion of hemocytes’ number after the placement of the artificial plug. Besides, the artificial plugs of *Z. fuscus* presented a minimal and homogeneous encapsulation. In contrast, females of plug producing species showed a decrease in the number of circulating hemocytes, accompanied by greater encapsulation on certain zones of the artificial plugs. These findings suggest that species with genital plugs (*Urophonius* spp.) have developed a more sensitive immune system to specific challenges like the one used in this study. In scorpions, the cuticle thickness and histological complexity of the female’s genital atrium have been related to mechanical damage caused by the capsular eversion of the spermatophore or by the introduction of genital plugs [12,18,84]. For instance, *Z. fuscus*, a non-plug producing species, presents simple spermatophores and a thin walled genital atrium, whereas plug producing bothriurid species have more complex spermatophores, and the genital atrium with folded epithelium and thick cuticular walls [18]. In addition, in several bothriurids, including *U. brachycentrus*, there have been seen regions in the atrium’s apical zone which contain epithelial cells with microvilli and pores connected to ducts [18,84]. We also found differences in the encapsulation response in the different zones of the artificial genital plug, especially in the plug producing species. There is evidence that female’s genitalia is complex and may present a modularized immune response (specificity) [47,119-121]. This specificity can be explained because the female’s tract has evolved by making contact with sperm, with male’s ejaculate substances and with infectious agents [120,122]. Preliminary data on *U. brachycentrus* suggests that the immune response triggered by artificial implants in females’ genitalia may be much more specific and intense than that triggered in other non-genital parts of their bodies. For example, there has been observed a weaker encapsulation response in implants placed in the dorsal pleural membrane compared to that triggered on artificial plugs in the genital area (Oviedo-Diego, Mattoni, Peretti personal observations).

### Types of identified hemocytes and quantified hemocyte load

We have described, for the first time, the types of hemocytes found in the female’s hemolymph in *Urophonius achalensis, U. brachycentrus* and *Z. fuscus*. Two main types of cells were found: plasmatocytes (PLs) and granulocytes (GRs), in agreement with the findings of existing works on the subject [69,102,123-126]. Subtypes of hemocytes, Cystocytes (CYs), Spherulocytes (SPs) and Adipohemocytes (ADs), previously cited for scorpions [69,102,123-124] were also identified. The existence of several types of hemocytes in the hemolymph would be an ancient character [127], which could have been retained in scorpions, one of the oldest arthropod groups [128-131, but see 132]. There were not found prohemocytes, described as stem cells with embryonic nature [69,102,123,126], probably due to their rapid conversion to other cell types [133]. Even though other subtypes of hemocytes, like oenocytoids or coagulocytes, were observed in scorpion species [69,102,123-124) there was not found any evidence of their presence in the species studied herein. It would be very useful to include novel techniques such as genetic markers or antibodies in future studies, as well as other microscopy techniques such as scanning or electronic transmission, for the purpose of a precise classification, quantification and elucidation of the action mechanisms of these cells [124,134-137]. *Zabius fuscus* had the highest values of THL compared those of plug producing species. The causes of this difference could respond to the evolutionary history of each species and the sexual and ecological context in which they have evolved [138-139]. Although the characteristics of the habitats are similar, the species present contrasting characteristics regarding patterns of surface activity at different times of the year [91-92]. In addition, these species could exhibit differences in microhabitats (Oviedo-Diego, Mattoni, Peretti personal observations), or in other parameters such as diet or potential parasites [83].

### Correlations between immune parameters

Since multiple immunological parameters can be costly to maintain [74] trade-offs may exist between parameters within the same system. This would indicate an overlap in the resources used by different defense mechanisms, a cross-regulation between them or a common underlying mechanism [64,61,138]. We found a negative correlation between the areas and the coloration of the encapsulations. This would suggest a trade-off between the encapsulation response per se (aggregation of layers of hemocytes) [140] and the melanization that occurs in these encapsulations (products generated by the phenoloxidase pathway) [66]. There was a close interrelationship between the humoral and cellular components [141], and some studies have reported antagonisms between the parameters of these systems, without investigating the underlying physiological mechanisms [97,142-143]. On the other hand, we found a positive correlation between body weight and THL. Variation in immunological parameters between individuals and species is expected [138,144]) since the management of trade-offs between the costs of immune defense and other life history traits that overlap in the use of resources, can vary (75,145]. Heavier individuals may be more immunocompetent since they would have more circulating hemocytes [146, but see 147]. However, it was also found that higher body weight individuals (*Z. fuscus* females) presented smaller areas of melanotic encapsulations on the artificial plug and clearer encapsulations. These results were expected since a higher THL value would indicate a higher concentration of free hemocytes in hemolymph, and a lower number of hemocytes in the genital area, resulting in less encapsulated and melanizated artificial plugs. It has also been reported that individuals with large numbers of hemocytes have a lower proportion of phagocytic hemocytes [148].

## Conclusions

The morphology of genital plugs of two scorpion species (*U. achalensis* and *U. brachycentrus*) was described, being complex structures with three different zones, strongly anchored to the female genital atrium, with a coloration that considerably resembles a type of immune response. The encapsulation and melanization patterns on the artificial plugs may indicate greater and more specific response in females of species that have a genital plug. In turn, these species presented a depletion in the number of hemocytes in hemolymph, indicating possible recruitment of these cells into the genital area. Body weight was correlated positively with THL but negatively with the encapsulation area on the artificial genital plug. These results and the negative correlations between different immunological parameters may indicate complex interrelations within the immune system that remains to be investigated. These results suggest that females of *U. achalensis* and *U. brachycentrus* contribute to the formation of the genital plug by attaching encapsulations to it and darkening the plug in certain zones by melanization of these encapsulations. Further studies would include analysis of changes in THL comparing virgin and inseminated females (with genital plug) to elucidate whether the observed results with artificial plugs actually reflect what happens when females are plugged. A comparison of the immune response triggered by implantation in the genital area with respect to other body regions and comparisons with the immunological parameters of the male would provide information on the specificity of the female’s genital immune response. Finally, modulation of female’s immune response according to male’s characteristics would provide information about female cryptic choice mechanisms, or about the female’s influence on the effectiveness of the plug to avoid sperm competition.

## Acknowledgments

We would like to thank J. Heywood for the revision of the language of the manuscript, to G. Gonzalez for his help with statistical questions, to D. Fink for collaborating in the artwork. Also, we are grateful to L.E. Costa-Schmidt and all members of LAB.R.E, especially P. Olivero and F. Bollatti for their useful suggestions on a previous version of the manuscript.

## Supporting information

**S1 Table. Mean values and standard deviations of different immunological parameters of *Urophonius brachycentrus*, *U. achalensis* and *Z. fuscus*.** The total hemocyte load (THL) before and after implantation of the artificial genital plug, and the decrease in hemocyte concentration between both stages are presented. The melanotic (ME) and non-melanotic (NME) encapsulation response was measured in all three zones of artificial genital plugs. a, b, and c indicate the grouping and separation between stages and zones (p <0.05) of the artificial genital plug. Capital letters (A,B) indicate the grouping and separation between species (p <0.05).

